# Systematic discovery and classification of human cell line essential genes

**DOI:** 10.1101/015412

**Authors:** Traver Hart, Megha Chandrashekhar, Michael Aregger, Zachary Steinhart, Kevin R. Brown, Stephane Angers, Jason Moffat

## Abstract

The study of gene essentiality in human cells is crucial for elucidating gene function and holds great potential for finding therapeutic targets for diseases such as cancer. Technological advances in genome editing using clustered regularly interspaced short palindromic repeats (CRISPR)-Cas9 systems have set the stage for identifying human cell line core and context-dependent essential genes. However, first generation negative selection screens using CRISPR technology demonstrate extreme variability across different cell lines. To advance the development of the catalogue of human core and context-dependent essential genes, we have developed an optimized, ultracomplex, genome-scale gRNA library of 176,500 guide RNAs targeting 17,661 genes and have applied it to negative and positive selection screens in a human cell line. Using an improved Bayesian analytical approach, we find CRISPR-based screens yield double to triple the number of essential genes than were previously observed using systematic RNA interference, including many genes at moderate expression levels that are largely refractory to RNAi methods. We further characterized four essential genes of unknown significance and found that they all likely exist in protein complexes with other essential genes. For example, RBM48 and ARMC7 are both essential nuclear proteins, strongly interact and are commonly amplified across major cancers. Our findings suggest the CRISPR-Cas9 system fundamentally alters the landscape for systematic reverse genetics in human cells for elucidating gene function, identifying disease genes, and uncovering therapeutic targets.

---

The Human Genome Project has yielded a fairly complete catalogue of cellular components, and a major goal moving forward will be to systematically discover and classify all genetic elements involved in normal biological processes and disease (*1*). With advances in gene editing technologies enabled by the CRISPR-Cas system (*2*–*5*), it is no longer quixotic to seek a comprehensive picture of cellular circuitry for human cells. Fortunately, yeast research has helped to establish a conceptual framework for systematic genetics, providing a glimpse of the basic modular organization of a cell and the corresponding genetic landscape (*6*). Much of the basis for advances in yeast systematic genetics was the discovery and classification of essential and non-essential genes (*7*, *8*). However, this binary classification was later demonstrated to be an artifact of laboratory conditions: while some genes do indeed seem to be constitutively essential, many others revealed fitness defects only under environmental stress or in different genetic backgrounds (*9*, *10*).

The distinction between core and context-dependent essentials is particularly relevant in humans, where a typical cell expresses perhaps two-thirds of the genome’s complement of genes (*11*). Identifying a tissue’s essential genes, and delineating them from the constitutive essentials shared across all tissues, may hold the key to unlocking tissue-specific disease. In tumors, this is the foundation for the concept of target identification by synthetic lethality (**12**): genes essential in tumor cells but not in adjacent normal tissues should make ideal therapeutic targets with high effectiveness and minimal side effects.

Identifying these context-specific essentials has been no easy feat. To date, this field has been constrained by technology; RNA interference (RNAi) has been the best available tool but, despite improvements in reagent design and analytical approaches, its utility is limited by imperfect mRNA knockdown as well as confounding off-target effects (*13*–*15*). We previously exploited the concept of core vs. context essentials – the Daisy model of gene essentiality, where each petal represents a tissue or context but all tissues share the core – in a systematic analysis of RNAi and first-generation CRISPR library screens (*16*). Our approach asserts that the degree to which a given screen recapitulates a known core essential set is a measure of its accuracy. One CRISPR screen showed marked improvement over comparable shRNA screens, hinting at the existence of a large pool of human essential genes that may be beyond the reach of RNAi. However, a screen in a second cell line by the same team yielded results only marginally better than random expectation, highlighting the wide variation in CRISPR efficiency and casting doubt on the generalizability of the system.

## Generation and testing of an ultracomplex human CRISPR library

We sought to confirm the utility of CRISPR as an improved screening technology to extend the catalogue of both the core and context human essential genes. We designed a genome-scale library of 174,978 guide RNAs (gRNAs) targeting 92,741 coding exons or adjacent splice sites in 17,661 protein-coding genes (see Supplementary Information). Over 70% of the gRNAs (126,900) have no predicted off-target binding site in hg19, even allowing for up to 2 mismatches, with the remainder (48,078) having only a single predicted 2-mismatch off-target site. We also added control sequences targeting non-human genes including EGFP, LacZ, and luciferase, as well as 1,380 gRNAs targeting random loci on chromosome 10, more than half of which are “promiscuous gRNAs” with ≥20 genomic binding sites (range = 20-5×10). The total library contains 176,500 gRNAs.

We established clones of HCT116 colorectal cancer cells that constitutively express *Streptococcus pyogenes* Cas9, and confirmed the editing ability of two independent clones (C1 and C2) following lentiviral-based expression of gRNAs targeting PTK7, TP53 and CEP85 (Fig. 1A, S1, S2A). These gRNAs decreased protein expression of target genes as measured by western blot analysis (Fig. 1B, Fig. S2B). RNA sequencing analysis of the parental HCT116 cell line, as well as the two Cas9-expressing clones, revealed no major gene expression changes (Fig. S3, R=0.99). We then tested whether gRNAs targeting essential genes caused anti-proliferative effects following expression in HCT116-C1 cells (hereafter referred to as “HCT116”). We cloned gRNAs targeting the essential genes PSMD1, EIF3D and PSMB2 into pLCKO from our own library designs, as well as from an independent design (Fig. 1C) (*17*). At 10 days post-puromycin selection, gRNAs targeting all three genes showed moderate to strong antiproliferative effects relative to the control LacZ gRNA (Fig. 1D; p<0.05, Student’s T-test). gRNAs we designed caused anti-proliferative effects comparable to (or better than) other designs (Fig. 1D, see starred gRNAs). Together, these results suggested that our HCT116 cells and gRNA designs were capable of identifying cancer cell proliferation genes.

**Figure 1.**
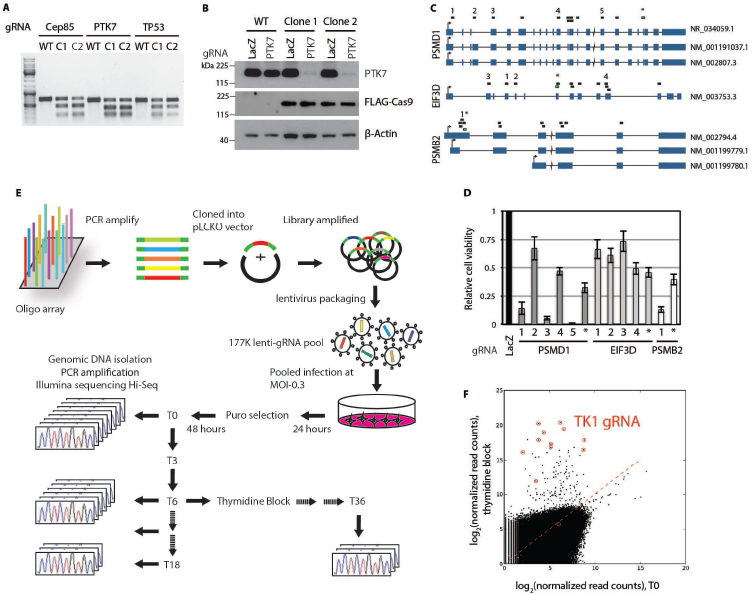
gRNA library testing and experimental setup of a genome-scale screen for essential genes in HCT116 cells. (A) Validation of the editing efficiency of the CRISPR-Cas9 system. Parental HCT116 cells (WT) and two single-cell derived HCT116 clones stably expressing FLAG-Cas9 (C1 and C2) were infected at low MOI with lentivirus harboring gRNA-expressing cassettes targeting CEP85, PTK7 or TP53. Infected cells were selected with puromycin for 48 hours, and 8 days post-selection Cas9 editing efficiency was monitored by the T7 endonuclease cleavage assay. (B) As in (A) but Cas9 editing efficiency was monitored by western blotting with antibodies against PTK7. Cas9 was detected with anti-FLAG antibodies and β-Actin was used as a loading control. (C) Gene models of essential genes including PSMD1, EIF3D, and PSMB2 showing target locations of library-designed gRNAs as small boxes above each model. The numbers and asterisks indicate the individual gRNAs used in part (D). (D) Cell viability assays in HCT116 cells infected with lentiviral-based gRNA expression cassettes. Target genes and gRNAs are indicated on the horizontal axis, and cell viability as measured by AlamarBlue staining 10 days post-selection relative to the LacZ control is indicated by the vertical axis (n=3). (E) Construction of the lentiviral pLCKO gRNA library and experimental setup for negative and positive selection screens. (F) Positive selection screen for suppressors of thymidine block. Normalized read counts of all gRNAs at T0 are plotted against the mean normalized count for thymidine-treated samples. TK1 gRNAs are circled in red.

We then cloned the 177k library into the pLCKO vector at 650-fold representation and confirmed this by deep sequencing. The plasmid library was converted to a lentiviral library pool, which was used to infect HCT116 cells at an MOI~0.3, or an average of one clone per cell (*18*), where each gRNA was represented at 270-fold coverage in the cell population (Fig. 1E).

To evaluate our library, we conducted a straightforward positive selection screen to look for suppressors of a cell cycle block. We performed the positive selection screen in HCT116 cells by applying a thymidine block 6 days after puromycin selection for cells infected with our gRNA library. The thymidine block was maintained for >4 weeks until clonal proliferation and outgrowth was observed (Fig. 1E). Genomic DNA from the outgrown cells was harvested and subjected to high-throughput sequencing, resulting in a strong enrichment of cells expressing gRNAs targeting thymidine kinase 1 (TK1). In fact, 11 out of 12 TK1 gRNAs present in the library were significantly enriched, while no other gene was represented by more than one gRNA (Fig. 1F). In the presence of high concentrations of thymidine, dTTP production is elevated causing an imbalance of dNTPs, which leads to cell cycle arrest at the G1-S transition through the activity of TK1 (Fig. S4 and(*19*)). Although TK1 has been previously identified as a factor in eliciting a thymidine block in mammalian cells (*20*–*22*), it has not been reported as directly identified in an unbiased genetic suppressor screen.

To identify essential genes, we performed parallel negative selection screens and sampled evolving cell populations every 3 days from day 6 to day 18, or approximately 20 doublings (Fig. 1E). Genomic DNA from each time point was isolated, and gRNA abundance was measured by massively parallel sequencing of the integrated gRNA cassettes in order to monitor the change in abundance of each gRNA between the initial cell population and each of the subsequent 5 time points (see Supplementary Information). gRNAs targeting essential genes are expected to “dropout” from the initial population, while those targeting non-essential genes are expected to be maintained (*18*, *23*).

The severity of phenotype that can be expected from off-target effects of a single gRNA is largely unknown. To address this question upfront, we analyzed the 827 gRNA sequences in our ultracomplex library that target random sites on chromosome 10, and that were detected with at least 50 reads at T0. Of these, 479 were designed using the same stringency filters as the gRNAs targeting coding exons; that is, none have predicted off-target binding sites within two mismatches. The remaining 348 gRNAs are “promiscuous gRNAs” with more than 20 or more predicted perfect match off-target binding sites. We measured the fold-change distribution of each set at each time point and observed that specific targeting of random sites on chr10 yields a fold-change distribution similar to that of gRNAs targeting nonessential genes, while promiscuous gRNAs induce a severe shift in fold-change distribution (Fig. S5). Whether this indicates that multiple genetic lesions are highly toxic or simply reflects the relationship between increased off-target sites and increased likelihood of cleaving a critical locus is unknown. Surprisingly, we do not observe a correlation between number of promiscuous target sites and population dropout rate (Fig. S6). Follow-up studies that measure the relationship between the number of predicted genomic target sites and the corresponding fitness defect could specifically assay the robustness of the genome and potentially improve the design of next-generation gRNA libraries by relaxing needlessly strict target specificity constraints.

## Analyzing negative selection screens

We adapted the analytical pipeline described previously in Hart *et al* to evaluate the quality of our dropout screen and to classify essential genes (*16*). Using the gold-standard sets of 360 essential and 927 nonessential genes defined in that study, we observed that the fold-change distribution of gRNAs targeting essential genes is significantly shifted relative to those targeting nonessential genes, and that the shift increases with time (Fig. 2A). We then applied an updated version of the algorithm described in (*16*), which implements dynamic boundary selection and 5-fold cross-validation to facilitate general applicability to negative selection screens. The improved version of this algorithm, which can easily be adapted for custom screening purposes, is referred to as the Bayesian Analysis of Gene Essentiality (“BAGEL”). We used BAGEL to calculate a log Bayes Factor (BF) for each gene (see Supplementary Information), with more positive scores representing higher confidence that a given gene’s knockout causes a decrease in fitness—for which we use the general term “essential”. Unlike most other analytical methods for large-scale gene perturbation studies, BAGEL uses the data from all reagents and all samples in a screen, and therefore provides a ready framework for the integration of experiments with multiple time points. Moreover, the use of gold-standard reference sets enables the unbiased evaluation of screen performance using precision and recall (Fig. 2B).

**Figure 2.**
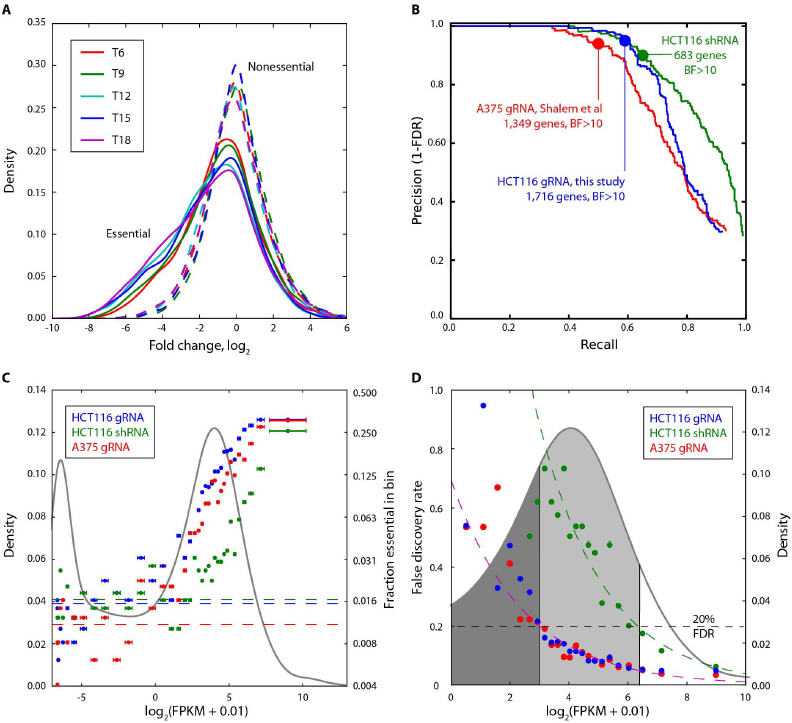
Evaluating screen performance. (A) Fold-change distributions from normalized gRNA read counts for gRNAs targeting gold-standard essential (solid) and nonessential (dashed) genes for each of the indicated time points. (B) Precision-recall curves after applying BAGEL to the HCT116-gRNA, HCT116-shRNA and A375-gRNA screens. (C) Quantile normalized gene expression density estimate for HCT116 and A375 (gray curve; left Y-axis). Genes are rank ordered by expression and binned (n=500). For each bin, mean expression (+/− s.d.) is plotted against the fraction of essential genes (BF>10; right Y-axis; note log scale). Mean %essential of bins with FPKM<-2 (dashed line) estimates background error rate of each screen. (D) Background error rate divided by the %essential is a binwise FDR estimate. Shading highlights CRISPR and RNAi effective regions.

We compare the performance of the HCT116 screen to a similar screen for essential genes in the A375 melanoma cell line (*24*), which we reanalyzed with the BAGEL pipeline, as well as a genome-scale pooled shRNA dropout screen we performed in HCT116 cells using a library of 78,432 shRNAs targeting 16,056 genes (*25*). Both CRISPR screens captured a large fraction of the reference essential gene set (~30% recall) without misclassifying any reference nonessentials; that is, a false discovery rate of 0%. (Fig. 2B). The shRNA screen has higher overall recall, presumably because the reference sets are derived from repeated observations in shRNA screens (*16*, *23*, *26*). However, the total number of genes classified as essential is markedly higher in CRISPR screens: 1,716 in HCT116 cells and 1,349 in A375 cells at a BF > 10, a threshold that identifies essential genes at high accuracy, compared to just 683 genes in parental HCT116 cells by shRNA (Fig. 2B).

We further evaluated screen quality by an independent method comparing essential genes to gene expression. Fig. 2C shows the distribution of log_2_(FPKM + 0.01) gene expression values (see Supplementary Information for details), with the main peak representing the bulk of active gene expression and the left peak representing unexpressed genes. We rank-ordered genes by expression level and divided them into bins of 500 genes; for each bin we then plotted the mean expression level versus the fraction of genes classified as essential in each screen (Fig. 2C). A background error rate for each screen was then calculated by taking all genes with trace/no expression (logFPKM < −2) and measuring the fraction classified as essential, assuming that these trace/no expression genes are false positives (Fig. 2C, dashed lines). At our threshold of BF>10, all three screens had a comparable background error rate of 1.1-1.6%, though the CRISPR screens clearly identified many more essential genes across all relevant gene expression levels (Fig. 2C). Moreover, our ultracomplex library detected 27% more essential genes than the 65K member GeCKO library used by Shalem *et al*. (*24*), across all expression levels.

The background error rate enables the analysis of how false discovery rates vary with gene expression. The binwise false discovery rates (the background error rate divided by the fraction of essential genes in each bin) for both the HCT116 and A375 CRISPR screens are virtually identical, with genes expressed at logFPKM > 3 having a binwise FDR < 20% (Fig. 2D). This compares very favorably against pooled-library shRNA screens, which achieve a comparable FDR above logFPKM 6.25. When projected onto a density plot of gene expression, the enormous advantage of CRISPR over RNAi is highlighted (Fig. 2D): over 8,100 genes are expressed in the effective CRISPR window, comprising some 70% of the ~11,600 genes with biologically relevant expression levels (logFPKM > 0), compared to just ~1,500 genes (13~) assayable by pooled-library shRNA with comparable accuracy. For CRISPR, the relationship between expression and error rate is likely driven by the fact that virtually all essential genes in a cell are expressed at these moderate to high levels, as there is no *a priori* reason why CRISPR, which targets DNA, should have an expression bias (contrary to RNAi, which targets transcripts).

The high accuracy of CRISPR screens analyzed by BAGEL across a wide range of expression levels is reflected in the concordance between the two CRISPR screens. Performed by different labs using different cell lines, libraries, and experimental designs, and using a fundamentally new technology, the two screens nevertheless detected 662 genes in common, or nearly 50% of the smaller Shalem *et al*. set. Of these genes, over 90% (n=601) are expressed at logFPKM > 3, where this intersection represents a 2.9-fold enrichment over random expectation (P = 1.4×10^−11^, hypergeometric test). Nevertheless, each screen captures a large number of unique essentials, and neither screen exceeds 60% recall against the gold-standard reference set (Fig. 2B) at the specified threshold, suggesting a substantial false negative rate and a large population of still-undetected essential genes. Intersection analysis estimates the total population of cell-line essentials at ~2,800 genes, which is comparable to the estimates from the precision-recall analysis for A375 (1,349 genes/50% recall = 2,698 genes) and HCT116 (1,716 genes/59% recall = 2,908 genes). These highly concordant estimates nearly triple the number of essential genes predicted by RNAi and represent a fundamental advance towards comprehensive reverse genetic screens in mammalian cell lines.

While the intersection between the two screens is a measure of their accuracy, the differences between them provide insight into real biological differences. Notably, knockout of mitochondrial genes causes fitness defects in HCT116 colorectal cancer cells but not in A375 melanoma cells (n=30 genes, 3.0-fold enrichment, P-value = 10^−7^), consistent with the BRAF-driven melanoma cell line’s dysfunction in oxidative phosphorylation and addiction to glycolysis (*27*). Conversely, all three members of the CDK-activating kinase (CAK) complex, including CCNH, CDK7, and MNAT1, are essential in A375 but not HCT116, suggesting a unique dependency of melanoma cells on CDK activity and a potential therapeutic window for existing CDK inhibitors.

## Characteristics of CRISPR essential genes

Functional genomics in model organisms and in human cell lines has proven that subunits of stable protein complexes tend to share the same loss-of-function phenotype (*28*). We tested the 1,716 HCT116 essentials for enrichment in CORUM and GO protein complexes, identifying 30 largely non-overlapping protein complexes that are highly enriched for essential subunits (Bonferroni-corrected P-value < 0.05; Fig. 3A). In addition, we compared the number of essential subunits in each complex to the number of subunits in the list of 823 global essentials gleaned from high-quality shRNA screens (*16*). In all cases, CRISPR increased the fraction of subunits classified as essential. Remarkably, over a third of these protein complexes are detected exclusively by CRISPR, where 0 or 1 subunits were detected by RNAi (Fig. 3A). Among the 121 essential subunits of these complexes, nearly 70% (n=83) have moderate mRNA expression (logFPKM from 3 - 6.25) and only 3 have lower expression (Fig. 3B). In most cases, these complexes perform basic biological processes such as DNA replication; for example, the origin recognition complex, the minichromosome maintenance complex, the DNA synthesome, kinetochore proteins, the centromere complex, the RNA polymerase I complex, the RNA polymerase III complex, and the transcription elongation complex (Fig. 3A). Of the 35 high-expression genes detected exclusively by CRISPR, 31 are mitochondrial, consisting of subunits of the mitochondrial ribosome and Complex I of the electron transport chain (Fig. 3A–B).

**Figure 3.**
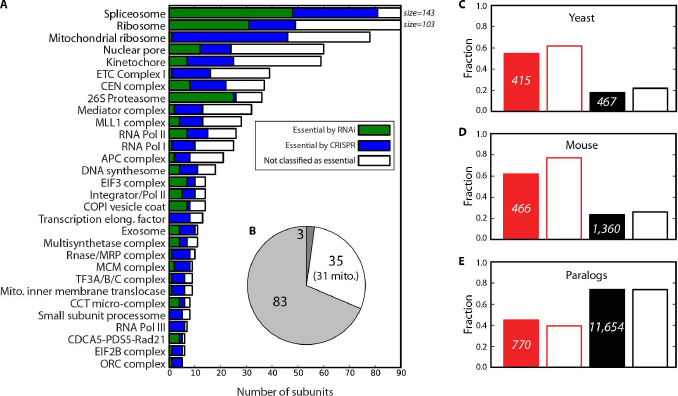
Characteristics of CRISPR essentials. (A) Selected essential protein complexes and detection rates by CRISPR (blue) and RNAi (green). (B) Genes detected by CRISPR but not RNAi, shaded by expression level (Fig. 2D). (C) Fraction of CRISPR essentials whose yeast orthologs are also essential (red, solid), compared to a strict set of “essentials in essential complexes” from RNAi (red border), CRISPR nonessentials (black, solid), and RNAi nonessentials (black border). (D) Fraction of CRISPR essentials whose mouse knockouts are prenatal lethal, as in (C). (E) Fraction of CRISPR essentials with one or more paralogs, as in (C).

Beyond enrichment for protein complexes, the CRISPR essentials exhibit genomic and evolutionary characteristics consistent with previous observations of essential genes. More than half of CRISPR essentials are also essential in yeast (Fig. 3C), and more than 60 percent have mouse orthologs whose knockouts cause prenatal lethality (Fig. 3D). These ratios are strikingly similar to those observed when comparing a much smaller set of 231 essential genes in essential complexes derived from RNAi (*16*). In addition, CRISPR essentials also show the same depletion for paralogs as RNAi essentials (Fig. 3E). The similarity of these signatures, despite the fact that the CRISPR set is more than sevenfold larger than the set of subunits of RNAi essential complexes, argues strongly for the accuracy of the set and the much greater sensitivity of the CRISPR screening assay.

## Characterizing novel essential genes

There still remains a significant portion of human genes that are uncharacterized or unstudied, part of the “dark matter” of the genome. We selected a subset of essential but uncharacterized genes from the HCT116 screen for further study. We chose eleven hits which each have less than the median number of Pubmed citations and GO terms for essential genes (Fig. 4A–B and Supplementary Information). The list was winnowed further to genes that are available in the ORFeome v8.1 cDNA collection, that were sequence verified, and that could be expressed in HEK293T cells, resulting in a final list of four genes (ANKRD49, ZNF830, CCDC84, RBM48). Notably, all four of these genes are expressed at moderate levels refractory to RNAi perturbation (Figure 4C). We confirmed that all four genes are essential for cell proliferation in HCT116 cells by independent treatment with single gRNAs. At 10 days post-puromycin selection, gRNAs targeting the genes caused moderate to strong anti-proliferative effects relative to controls (Fig. 4D, p<0.05, Student’s T-test).

**Figure 4.**
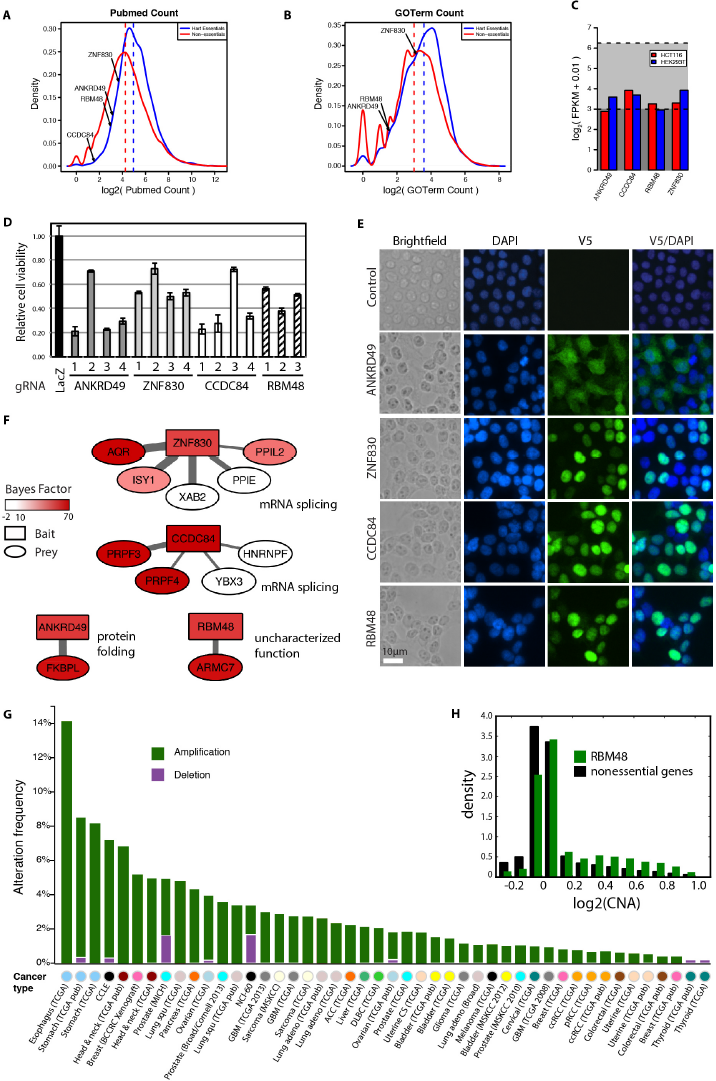
Validation of uncharacterized essential genes. Distributions of the number of Pubmed citations (A) or GO terms (B) linked to each gene symbol in Entrez Gene for genes found to be essential by CRISPR (blue), or non-essential (red). Dotted lines indicate median value for each distribution. Position of selected uncharacterized essentials in the distribution is shown. CCDC84 had no associated GO terms. (C) RNAseq expression level of the uncharacterized genes in both HCT116 cells (red) and HEK293T cells (blue). Dotted black lines denote the effective regions for CRISPR (lower line) and RNAi (upper line). (D) Cell viability assays in HCT116 cells infected with lentiviral-based gRNA expression cassettes, as measured by AlamarBlue staining 10 days post-selection (n=3). (E) Immunofluorescence microscopy analysis of 293T cells stably expressing V5-tagged ORFs. Subcellular localization of ORFs was detected with an anti-V5 antibody (green), and nuclei were stained using DAPI (blue). (F) Summary of the protein-interaction network of ANKRD49, ZNF830, CCDC84 and RBM48 detected by AP-MS analysis. V5- or 3xFLAG-tagged ORFs of the indicated proteins were immunoprecipitated from HEK293T and HCT116 cells, and interactors were identified by mass spectrometry analysis. Bait proteins are indicated by rectangles and prey proteins by circles. The color of the node corresponds to the Bayes Factor essentiality score, and the width of the edges corresponds to the number of times the interaction was detected. Results from 5 independent AP-MS experiments are shown for ANKRD49 and ZNF830, and 4 independent AP-MS experiments for CCDC84 and RBM48. (G) Cross-cancer alteration summary for RBM48. The histogram depicts the alteration frequency of RBM48 across tumor samples and cancer cell lines. The following alteration types are indicated: deletion (purple), amplification (green). This figure is modified from cBioPortal for Cancer Genomics {Cerami, 2012 #3498;Gao, 2013 #3499}. (H) Distribution of copy number across all tumors in cBioportal (n=11,545) for non-essential genes (black) and RBM48 (green).

We aimed to infer the function of each gene based on protein-protein interactions and subcellular localization. We expressed V5-tagged open reading frames (ORFs) of the four genes in HEK293T and HCT116 cells. ANKRD49 showed staining distributed throughout the cell, whereas ZNF830, CCDC84 and RBM48 exhibited exclusive nuclear staining (Fig. 4E and data not shown). These results indicate the subcellular locations where these proteins elicit their essential functions.

Next, we performed immunoprecipitations in HEK293T and HCT116 cells expressing V5-and 3xFLAG-tagged ORFs and analyzed the captured proteins by mass spectrometry (Fig. 4F and Supplementary Information). Interestingly, protein-protein interactions with other essential genes were detected for all ORFs tested, suggesting that these novel essential genes may also be assigned to essential complexes. For example, ANKRD49 was found to associate strongly with FKBPL, an essential gene in HCT116 cells with BF>20, and together likely function as an Hsp90 co-chaperone to promote aspects of protein folding (*29*). ZNF830 and CCDC84 reproducibly pulled down different sets of proteins that are predicted to participate in mRNA splicing (Fig. 4F). ZNF830 was previously found to associate with the XAB2 splicing complex, whereas CCDC84 has not been reported to interact with any splicing factors (*30*). Finally, the RNA-binding motif containing protein RBM48 captured ARMC7 as a prey, which itself was identified as a novel essential gene of unknown function. Interestingly, both RBM48 and ARMC7 are found to be amplified across several cancer tissues and cell lines (Fig. 4G, Fig. S7) (*31*, *32*). Across all tumors with copy number data in cBioPortal (n=11,545), both genes show an enrichment for copy number amplification and a corresponding depletion for copy loss relative to a pool of nonessential genes. (Fig. 4H). Collectively, these observations point to a novel, stable protein complex with no known function that likely has an RNA-related essential role in proliferating of tumor cells.

## Discussion

The adaptation of library RNA interference methods to mammalian cells a decade ago opened the door to large scale screens for synthetic lethal partners of known oncogenes. Though a great deal has been learned, the inherent limitations of RNAi and the research community’s failure to rigorously evaluate and standardize both experimental and analytical methods have collectively fallen short of the payoff that systematic reverse genetic screens have provided in model organisms. Now, with the emergence of the CRISPR-Cas9 system as a tool for genetic manipulation, expectations are again justifiably elevated. CRISPR library screens strongly indicate the existence of a population of essential genes perhaps three times larger than that predicted by RNAi, with an as-yet completely unknown breakdown between constitutive and context essentials. This fundamentally alters the landscape of systematic cancer target identification. New context-specific essentials represent potential synthetic lethal partners of known oncogenes, and new constitutive essentials broaden the net for genes whose haploinsufficiency, resulting from copy loss induced by genomic instability, may offer a therapeutic window for inhibition.

Though the potential is enormous, many of the same pitfalls that beset RNAi now lay in the path of widespread CRISPR adoption. Despite rapid progress in characterizing the CRISPR system, the community is largely unaware of what systemic shortcomings may exist - and exist they surely do. We offer an analytical pipeline that can, in principle, be applied to any large-scale library screen. Perhaps more importantly, we describe an evaluation framework that can be used to compare and improve future analytical approaches, and ideally prevent the false starts that have limited RNAi. With a foundation of rigorous analytical methods, the CRISPR-Cas9 system may finally enable systematic genetic research in mammalian cell lines on a scale to rival that achieved in model organisms.

## Acknowledgments

We thank all members of the Moffat and Angers labs; Dax Torti and the Donnelly Sequencing Centre for assistance with sequencing. This work was supported by grants to J.M. from the Ontario Research Fund, Ontario Institute for Cancer Research, Canadian Foundation for Innovation, and Canada Research Chairs program. J.M. is a Tier 2 Canada Research Chair and a Senior Fellow at the Canadian Institute for Advanced Research.

## Supplementary Information

Materials and Methods, Figures S1-S11, and Tables S1-S6 are not included in this preprint.

